# EEG functional network topology is associated with disability in patients with amyotrophic lateral sclerosis

**DOI:** 10.1101/065714

**Authors:** Matteo Fraschini, Matteo Demuru, Arjan Hillebrand, Lorenza Cuccu, Silvia Porcu, Francesca Di Stefano, Monica Puligheddu, Gianluca Floris, Giuseppe Borghero, Francesco Marrosu

## Abstract

Amyotrophic Lateral Sclerosis (ALS) is one of the most severe neurodegenerative diseases, which is known to affect upper and lower motor neurons. In contrast to the classical tenet that ALS represents the outcome of extensive and progressive impairment of a fixed set of motor connections, recent neuroimaging findings suggest that the disease spreads along vast non-motor connections. Here, we hypothesised that functional network topology is perturbed in ALS, and that this reorganisation is associated with disability. We tested this hypothesis in 21 patients affected by ALS at several stages of impairment using resting-state electroencephalography (EEG) and compared the results to 16 age-matched healthy controls. We estimated functional connectivity using the Phase Lag Index (PLI), and characterized the network topology using the minimum spanning tree (MST). We found a significant difference between groups in terms of MST dissimilarity and MST leaf fraction in the beta band. Moreover, some MST parameters (leaf, hierarchy and kappa) significantly correlated with disability. These findings suggest that the topology of resting-state functional networks in ALS is affected by the disease in relation to disability. EEG network analysis may be of help in monitoring and evaluating the clinical status of ALS patients.

## Introduction

Amyotrophic Lateral Sclerosis (ALS) is one of the most severe neurodegenerative diseases, affecting the upper and lower motor neurons. All motor functions are progressively invalidated, and life expectancy rarely exceeds 3 years from the onset of symptoms. However, in contrast to the classical tenet that ALS represents the outcome of extensive and progressive impairment of a fixed set of motor connections, recent neuroimaging findings suggest that the disease spreads along vast non-motor connections. Indeed, advanced neuroimaging techniques, which allow for the non-invasive investigation of structural and functional brain organization, have so far introduced new opportunities for the study of ALS and are currently supporting the multi-systemic pathophysiology of this disease^1^,^2^.

Recently, modern network science has aided in the understanding of the human brain as a complex systems of interacting units^3^,^4^. Indeed, the organization of brain networks can be characterised by means of several metrics that allow to estimate functional integration and segregation, quantify centrality of brain regions, and test resilience to insult^5^. Moreover, changes in network topology have been described for a range of neurological and psychiatric disorders^4^,^6^. In this view, structural and functional network studies based on diffusion tensor imaging (DTI) and functional magnetic resonance (fMRI) have contributed in elucidating basic mechanisms related to ALS onset, spread and progression.

For instance, Verstraete et al.^7^ observed structural motor network degeneration and suggested a spread of disease along functional connections of the motor network. Moreover, the same group has also reported^8^ an increasing loss of network structure in patients with ALS, with the network of impaired connectivity expanding over time. Schmidt et al.^9^, have recently shown that structural and functional connectivity degeneration in ALS are coupled and that the pathogenic process strongly affects both structural and functional network organization. Other resting-state fMRI studies^10–12^ have reported alterations in specific resting-state networks. Recently, Iyer and colleagues^13^ have investigated the use of resting-state electroencephalographic (EEG) as a potential biomarker for ALS, suggesting that a pathologic disruption of the network can be observed in early stages of the disease. However, it still remains relevant to address methodological issues that may affect both connectivity estimation and network reconstruction^14^.

Although the results described above are promising, it is not yet clearly understood how whole-brain functional networks are perturbed in ALS patients, and how this relates to disability. Resting-state EEG analysis may represent a practical tool to evaluate and monitor the progression of the disease. Despite the wide use of EEG in the assessment of brain disorders^4,15,16,^ it has not been used widely to evaluate functional network changes induced by ALS. To test our hypothesis, we reconstructed functional networks from resting-state EEG recordings in 21 ALS patients and 16 age-matched healthy controls using the phase lag index (PLI)^17^, a widely used and robust measure of phase synchronization that is relatively insensitive to the effects of volume conduction. The topologies of frequency specific minimum spanning trees (MSTs) were subsequently characterised and compared between groups as it has been shown^18,19^ that it avoids important methodological biases that would otherwise limit a meaningful comparison between the groups^20^. Moreover, a correlation analysis was performed between the MST parameters and disability.

## Results and Discussion

### Age-matching

No significant group differences were observed in age (W=145.5, *p* = 0.499).

### Functional Connectivity

No significant group differences were observed for the global mean PLI in any frequency band (both with and without FDR correction for number of frequency bands). Descriptive results and statistics are summarized in Table 1. No significant correlation was observed between the patients’ global mean PLI and the disability score for any frequency band (see Table 2).

**Table 1.** Group descriptive and statistics from Mann-Whitney U test for the global mean PLI.

| mean PLI | Group | N | mean | SD | MW | p-value | Cohen's d |
| --- | --- | --- | --- | --- | --- | --- | --- |
| delta PLI | ALS | 21 | 0.170 | 0.017 | 170.0 | .964 | 1.057 |
|  | Controls | 16 | 0.167 | 0.014 |  |  |  |
| theta PLI | ALS | 21 | 0.144 | 0.034 | 185.0 | .617 | 1.624 |
|  | Controls | 16 | 0.137 | 0.012 |  |  |  |
| alpha PLI | ALS | 21 | 0.153 | 0.041 | 165.0 | .940 | -0.396 |
|  | Controls | 16 | 0.155 | 0.043 |  |  |  |
| beta PLI | ALS | 21 | 0.070 | 0.008 | 113.0 | .095 | -2.589 |
|  | Controls | 16 | 0.073 | 0.007 |  |  |  |

**Table 2.** Correlations between global mean PLI and disability score.

|  |  | Disability score |
| --- | --- | --- |
| delta PLI | Spearman's rho | .021 |
|  | p-value | .929 |
| theta PLI | Spearman's rho | -.218 |
|  | p-value | .343 |
| alpha PLI | Spearman's rho | -.198 |
|  | p-value | .389 |
| beta PLI | Spearman's rho | -.351 |
|  | p-value | .119 |

### MST dissimilarity

A significant MST dissimilarity between ALS patients and healthy controls was found in the beta band using Mann-Whitney U test (W = 68.00, *p* = .008) after FRD correction.

### MST topology

A significant difference between groups was observed for MST leaf fraction in the beta band (W = 87.5, *p* = .014). Results from Mann-Whitney U test statistics are summarized in Table 3. Individual values for each MST parameter in the beta band are shown in Figure 1. In contrast, significant correlations were observed between some MST parameters (leaf, hierarchy and kappa) and disability score in the beta band (scatterplots for individual MST parameters and disability scores are reported in Figure 2).

**Table 3.** Group differences in the beta band.

| Beta band |  |  |  |  |  |
| --- | --- | --- | --- | --- | --- |
|  | W | p-value | mean differences | SE differences | Cohen's d |
| leaf | 87.50 | .014 | -0.033 | 0.012 | -5.202 |
| diameter | 186.00 | .591 | 0.005 | 0.009 | 2.027 |
| hierarchy | 104.00 | .051 | -0.017 | 0.009 | -3.730 |
| kappa | 111.00 | .083 | -0.299 | 0.189 | -3.405 |

**Figure 1.**
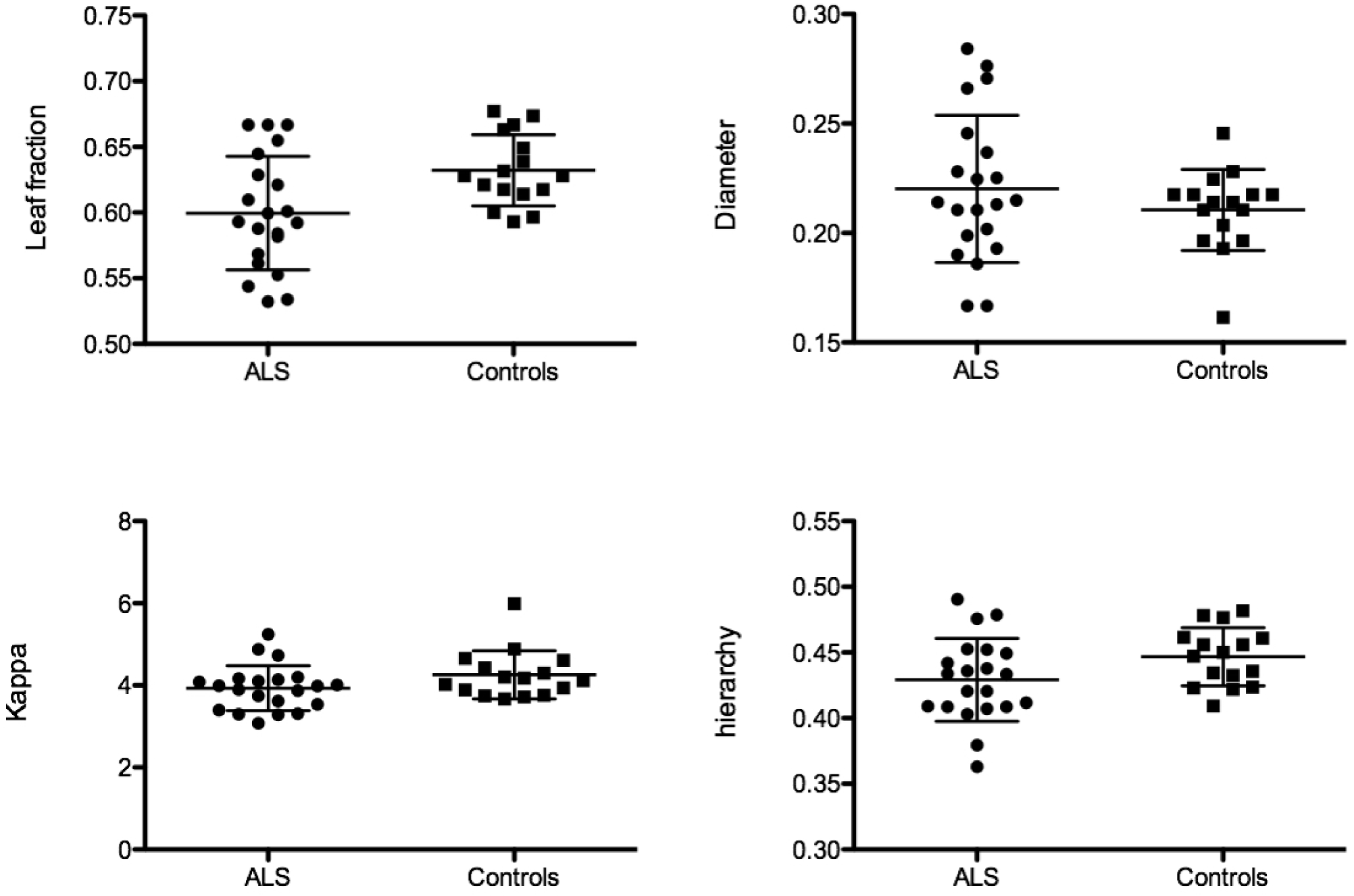
MST parameters for the patients and controls in the beta band. Horizontal bars indicate mean and standard deviation. Each dot or square represents a single ALS patients or healthy control, respectively.

**Figure 2.**
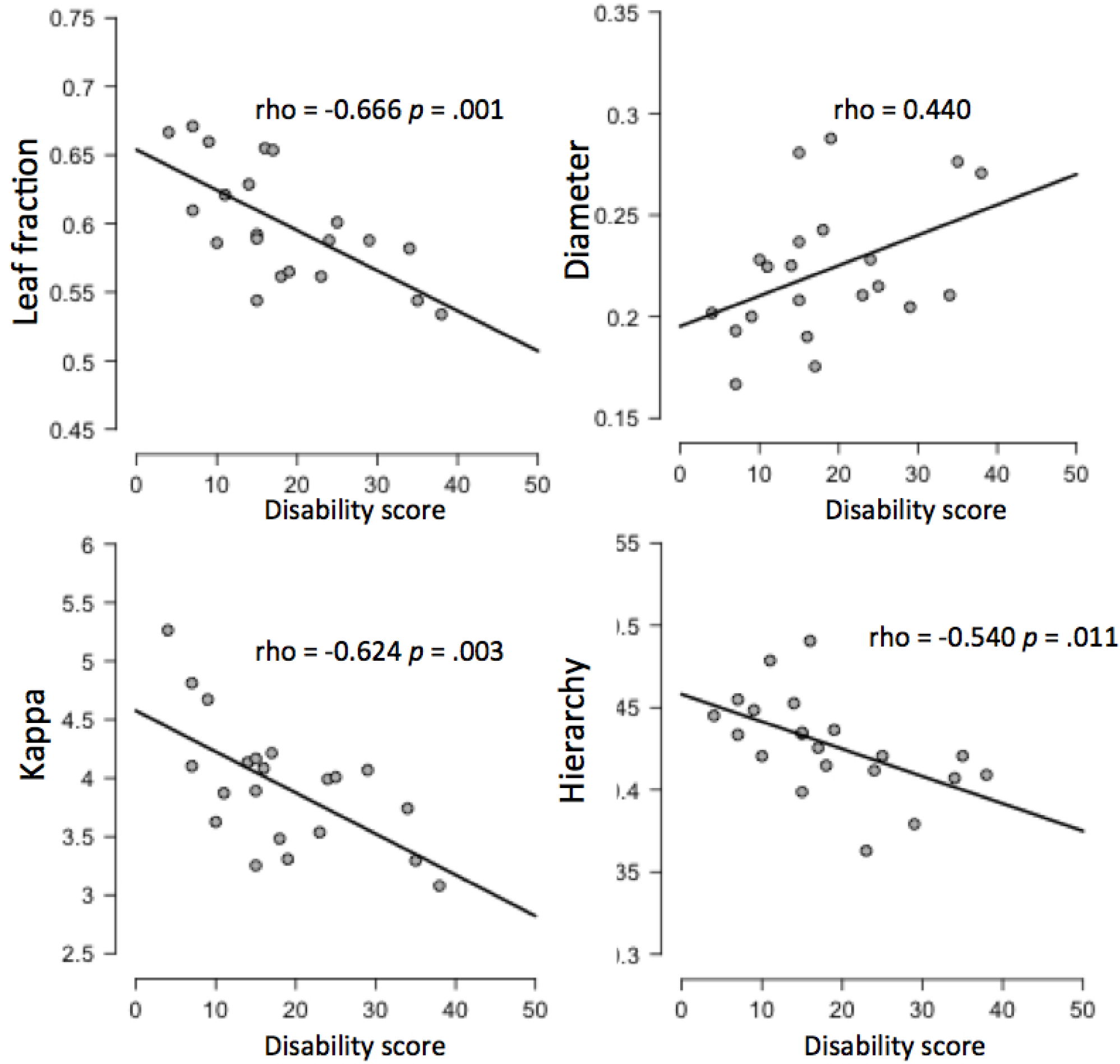
Scatter plots for MST parameters versus disability score in the beta band. Disability score was computed as (48 – ALSFRS-R), thus higher scores refer to higher disability.

**Figure 3.**
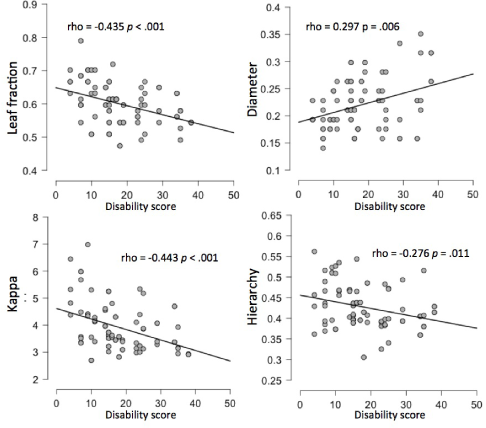
Scatter plots for MST parameters versus disability score in the beta band at single epoch level. Disability score was computed as (48 – ALSFRS-R), thus higher scores refer to higher disability.

### Discussion

In summary, by applying the PLI and the MST analysis in EEG recordings, this study shows large-scale changes in the functional brain network organization in ALS patients as identified using MST dissimilarity. Post-hoc analysis revealed that this difference in network topology between patients and controls was due to a difference in leaf fraction, and that the patients’ network organization in terms of MST parameters significantly correlated with disability, which is of clinical relevance. These results were observed in the beta band (13 – 30 Hz), where the MST topology was characterized by a significantly lower leaf fraction in the patients. Together with the significant negative correlation between disability score and leaf fraction, as well as a positive correlation with the diameter (even though not significant), indicates the tendency to deviate from a more centralized (star-like topology) towards a more decentralized organization (line-like topology). The negative correlation between tree hierarchy and disability score suggests that there is a sub-optimal balance between hub overload and functional integration in the network. Moreover, the negative correlation between disability and kappa, a measure that captures the broadness of the degree distribution, reflects the detrimental effect of a network topology with a reduced ease of synchronization (i.e., decreased spread of information across the tree)^21^.

As hypothesized, these findings suggest that ALS alters the brain network topology, which thus tends to deviate from the normal, presumably optimal, organization, and suggest that the correlation between MST parameters and the ASLFRS-R scale maybe be useful in monitoring the progression of the disease. These findings indicate that also at macroscopic scale (as measured by EEG functional networks), in accordance with previous studies on structural and functional neuroimaging^1^, ALS seems to affect extramotor brain regions, a result that is in line with the idea that pathological perturbations are rarely confined to a single locus^22^.

Moreover, it is of interest to note that a similar shift towards a more decentralized topology has been previously observed in multiple sclerosis^23^ and Parkinson’s disease^24^, and that functional networks in epilepsy patients that respond to vagal nerve stimulation re-organize towards a more centralized topology^25^. Together, these findings suggest that there is a possible common pathway in neurological disorders, as has been hypothesised recently^4^. Group differences and significant correlations between network topology and disability were found in the beta band, which may not be surprising given its link with motor function^26^ and that changes in beta activity can occur with ageing, sensorimotor disorders and amyotrophic lateral sclerosis^27,28^.

Despite the observed differences between healthy controls and ALS patients in terms of MST dissimilarity and leaf fraction, and clinically relevant correlations between disability and network topology, we found that the detection of distinctive EEG network properties still remains a difficult task during the early stages of the disease. This is in contrast with the study by Iyer and colleagues^13^, who used a set of connectivity metrics in combination with network analysis. In contrast to their work, we used different methods of functional connectivity (PLI) and network reconstruction (MST), as well as a conservative statistical approach. Indeed, the PLI has been shown to be an index of phase synchronization that is robust to biases introduced by volume conduction and field spread. Moreover, the MST represents a network approach that, although still providing network characteristics that can be related to conventional graph measures^19^, is not biased by common methodological issues arising when reconstructing and comparing networks using traditional approaches (i.e. the use of arbitrary thresholds)^20^.

Previous studies have shown a direct relation between disability and disease progression^29^. Interestingly, the observed correlation between network organization and disease disability suggests that it might be possible to track disease progression on the basis of EEG network analysis. However, a longitudinal study is needed to confirm this idea. The difference between groups in terms of overall MST topology (i.e. the MST dissimilarity results), in combination with the change in leaf fraction, suggests that ALS affects the brain networks at a global level. However, it could be that the networks were reconstructed at a level that was too coarse, and/or without enough anatomical precision. The study of source-reconstructed time-series would be of help in investigating the role of specific brain regions, and in identifying whether certain regions are more affected than others.

In conclusion, this study shows that EEG functional network re-organization in ALS patients, as computed by the PLI and MST approach, is associated with the patient disability. This finding suggests that resting-state EEG networks analysis may play an important role in evaluating the status of ALS patients and monitoring disease progression.

## Methods

### Subjects

Twenty-one patients (7 female; mean age 66, standard deviation 9 years) diagnosed with ALS according to the revised El Escorial criteria^30^, who attended the ALS Centre of the Azienda Ospedaliera Universitaria of Cagliari (Italy), were included in the study. A control group, consisting of sixteen age-and gender-matched healthy subjects (9 female; mean age 65, standard deviation 7 years), was also included. The local Ethical Committee approved the study (NP/2013/1496) and written informed consent was obtained from the participants. The clinical ALSFRS-R score^29^, a validated rating instrument for monitoring the progression of disability in patients with ALS, was evaluated at the time of EEG recording. This score was converted to a disability score by subtracting it from the maximum obtainable score, i.e. 48 - ALSFRS-R. That is, a disability score of 0 means that you are healthy.

### Recordings

Five minutes EEG signals were recorded using a 61 EEG channels system (Brain QuickSystem, Micromed, Italy) during an eye-closed resting-state condition. The reference electrode was placed in close proximity of the electrode POz. Signals were digitized with a sampling frequency of 256 Hz and offline re-referenced to the common average reference (excluding channels Fp1, AF3, AF7, Fp2, AF4 and AF8). For each subject the first four epochs (avoiding as possible contaminations from eye blinks, eye-movements, muscle activity, ECG, as well as systems-and environmental artifacts) of 2048 samples (8 s)^31^ were selected and band-pass filtered in the classical frequency bands: delta (1–4 Hz), theta (4–8 Hz), alpha (8–13 Hz), and beta (13–30 Hz).

### Functional connectivity

The phase lag index (PLI)^17^, which evaluates the asymmetry of the distribution of instantaneous phase differences between pairs of channels, was used to estimate functional connectivity (FC). Computing FC between all pair wise combinations of EEG time-series resulted in a weighted adjacency matrix of 58 x 58 entries (after excluding bad channels form both patients and healthy subjects) for each epoch. Mean PLI was also computed across epochs and channels. MST reconstruction. The MST is an acyclic sub-network, which connects all nodes minimizing the link weights (for the computation of the MST, the link weight is defined as 1 – PLI). The MST was obtained using the Kruskal algorithm^32^. The topology of the MST was characterised using several measures. The diameter (largest distance between any two nodes), the normalized leaf fraction (number of nodes with degree of 1 divided by the total number of nodes), kappa (broadness of degree distribution) and the tree hierarchy (balance between hub overload and network integration) were extracted from the MSTs^18,19, 33^. The procedure was repeated for each subject, each epoch and each frequency band separately.

### MST dissimilarity

MST dissimilarity, which assesses the overlap between MSTs^34^, was estimated between MSTs of ALS patients and healthy controls. In this study the MST reconstructed from the average connectivity matrix of all healthy subjects was used as reference in order to compute MST dissimilarities for both patients and controls^23^.

### Statistical analysis

Statistical differences in age, mean PLI, and MST dissimilarity between groups were evaluated using the non-parametric Mann-Whitney test. The value used for significance was set to p < .05 and a correction for multiple comparisons was performed by the false detection rate (FDR), correcting for the four frequency bands^35^. In case we found significant MST dissimilarity, post-hoc analysis was performed to find out which MST parameters were different. Moreover, a Spearman’s rank correlation coefficient was computed to assess the relationship between the network topology (in terms of mean PLI and MST parameters) and disease severity (in terms of the disability score). Statistical analysis was performed using JASP (version 0.7.5 beta 2 for Mac OS X)^36^.

## Acknowledgements

M. Fraschini and M. Demuru were in part supported by the Fondazione Banco di Sardegna (Prot.U7989.2013/AI.713.MGB. Prat.2013.1237).

## Author contributions statement

M.F., A. H., G.F., G.B. and F.M. conceived the study, S.P., F.D., M.P., M.D. and M.F. conducted the experiment, M.D., L.C. and M.F. analysed the data, A.H., M.F. and F.M. wrote the draft of the paper. All authors reviewed the manuscript.

## Additional information

The authors declare that they have no conflict of interest.

